# Dyskinesia induced by a dopamine 2 receptor-like agonist, but not a dopamine 1 receptor-like agonist, is blocked by sub-anesthetic ketamine

**DOI:** 10.64898/2026.10.27.762361

**Authors:** Raveena Parmar, Jasmine R. Meredith, Mitchell J. Bartlett, Emma G.E. Smith, Jeanelle M. Castro, Scott J. Sherman, Gaurav Chattree, Kathy Steece-Collier, Torsten Falk

## Abstract

L-DOPA remains the most effective symptomatic treatment for Parkinson’s disease, but chronic administration is frequently complicated by L-DOPA-induced dyskinesia (LID). Our prior work has established that sub-anesthetic ketamine can reduced LID acutely and has a long-lasting neuroplastic effect. This study tested whether low-dose ketamine reduces dyskinesia induced specifically by dopamine D2R-like or D1R-like receptor activation. Male Sprague Dawley rats were primed with daily L-DOPA to establish dyskinesia and then challenged with quinpirole, a D2R-like receptor agonist, or SKF-81297, a D1R-like receptor agonist, to induce abnormal involuntary movements (AIMs). After stable AIM baselines were established, animals received agonist plus vehicle or agonist plus ketamine. Limb, axial, and orolingual AIMs were scored on the acute treatment day (D0) and after washout on D4 and D7. Quinpirole and SKF-81297 reliably produced stable total LAO AIM scores following L-DOPA priming. Ketamine significantly reduced only quinpirole-induced, but not SKF-81297-induced, total AIMs compared with vehicle on D0, demonstrating an acute anti-dyskinetic effect. This reduction persisted after washout, with ketamine-treated animals continuing to show significantly lower AIMs on D4 and D7. Time-course analyses further showed reduced AIMs expression across the post-quinpirole observation period on D0 and at multiple time points on D7. These findings demonstrate that ketamine suppresses quinpirole-induced dyskinesia in L-DOPA-primed male rats and suggest that D2R-like receptor-mediated or indirect pathway-related mechanisms, but not direct pathway-related mechanisms, may contribute to ketamine’s anti-dyskinetic effects. Together, these results extend prior work on ketamine by linking its sustained anti-dyskinetic effects to dyskinesia driven by D2R-like receptor stimulation *in vivo*.

**Highlights:**

- Ketamine reduced quinpirole-induced abnormal induced movements on treatment day
- Ketamine’s anti-dyskinetic effect persisted after washout for 1 week
- D2R-like dopamine receptor-induced dyskinesia was sensitive to ketamine treatment
- D1R-like dopamine receptor-induced dyskinesia was not sensitive to ketamine treatment
- The findings suggest a possible indirect pathway involvement in effects of ketamine

## Introduction

L-DOPA-induced dyskinesia (LID) reflects abnormal dopaminergic stimulation within a dopamine-denervated striatum and is associated with maladaptive molecular, synaptic, and circuit-level plasticity in basal ganglia pathways [1,2]. Although D1-like receptor (D1R)-mediated mechanisms are often emphasized as the main driver of LID, given the importance of the direct or GO pathway, D2R-like receptor signaling is also important for understanding dyskinesia expression, particularly because D2 receptor-expressing striatal projection neurons (MSNs) are closely associated with the indirect or NOGO pathway [3,4]. And recent work indicates that ensembles of both behavior-encoding D1R- and D2R-expressing MSNs form new combinations of hyperactive neurons mediating specific dyskinetic features, and a D2R-specific pathway to LID was described by the Cenci and Colleagues [5,6]. The present work focuses on whether low-dose ketamine can reduce dyskinesia induced by D2R-like or D1R-like receptor activation and whether these findings support a possible role for indirect pathway-related or direct pathway-related mechanisms in the established anti-dyskinetic effects of ketamine [7–10].

The unilateral 6-hydroxydopamine (6-OHDA) lesion model provides a well-established platform for studying dyskinesia because chronic L-DOPA exposure produces reproducible AIMs in dopamine-denervated animals [11–13]. In this model, AIMs ratings, rotational behavior, and forelimb use have been pharmacologically validated as behavioral measures of dyskinesia and Parkinsonian motor impairment [12–14]. Limb Axial and Orolingual (LAO) AIMs scoring is therefore commonly used as a primary behavioral outcome for evaluating candidate anti-dyskinetic therapies in preclinical studies [12–14].

A major framework for interpreting dyskinesia involves the organization of striatal output pathways. Gerfen and colleagues demonstrated that D1R-related gene expression is associated primarily with striatonigral neurons, whereas D2R-related gene expression is associated primarily with striatopallidal neurons [3]. Later work refined this model by showing that dopamine regulates excitability, synaptic integration, and plasticity across D1R- and D2R-expressing striatal projection neurons in a dynamic and cell-type-specific manner [4]. This framework is important for dyskinesia research because chronic L-DOPA treatment does not simply restore normal dopamine signaling, but can instead produce abnormal stimulation of sensitized striatal circuits [2,4]. However, pharmacological activation of dopamine receptor families does not directly isolate a single neuronal population *in vivo*, so agonist-induced AIMs studies should be interpreted as receptor-linked behavioral probes rather than direct measurements of isolated direct or indirect pathway activity [4,15].

D1R-associated mechanisms have a prominent role in many models of dyskinesia, but D2R-like receptor activation can also contribute to abnormal motor output after dopaminergic priming. Delfino and colleagues showed that the unselective dopamine agonist apomorphine and the D1/D5 receptor agonist SKF-81297 induced strong forelimb dyskinesia and axial dystonia, whereas the D2R-family agonist quinpirole induced milder dyskinesia with chronic treatment [16]. These findings support a major role for D1/D5 receptor stimulation in peak-dose dyskinesia while also demonstrating that D2R-family receptor stimulation can elicit dyskinetic behavior under primed conditions [16]. More recent studies have further implicated D2R-expressing neurons and striatopallidal circuitry in LID expression and modulation, supporting continued investigation of D2R-like receptor-linked mechanisms in dyskinesia [17–20]. Because the indirect pathway is strongly influenced by D2R-expressing striatal projection neurons, pharmacological stimulation with quinpirole provides a useful approach for testing whether dyskinetic behavior can be elicited through D2R-like receptor activation after L-DOPA priming [15,16]. Generally, abnormal information storage in corticostriatal synapses is linked with the development of L-DOPA-induced dyskinesia [22].

Glutamatergic signaling is another important contributor to dyskinesia-related plasticity. Prior work from our group directly examined whether NMDA receptor antagonism differentially affects dopamine receptor agonist-induced AIMs. While earlier work from Paquette et al. showed that the highly selective antagonist MK-801 is only anti-dyskinetic at doses that are pro-parkinsonian [23], Flores and colleagues showed that MK-801 had differential effects on D1R- and D2R-induced AIMs, supporting the idea that glutamatergic modulation can interact differently with dopamine receptor-linked dyskinesia pathways [15]. Amantadine, another multimodal drug that has NMDAR antagonist activity, was recently shown to differentially change D1R/D2R neural ensembles in the striatum of rodent model of PD and LID [24]. Ketamine is relevant because it is also a multimodal ligand with noncompetitive NMDA receptor antagonist activity with additional pharmacological actions that may influence glutamatergic and opioid transmission, synaptic plasticity, oscillatory activity, and neuromodulatory systems [8–10; 25–27]. In addition, ketamine is also a positive allosteric modulator for opioid receptors [28]. Our previous work demonstrated that sub-anesthetic ketamine reduces LID in the 6-OHDA rat model and that a 10-hour low-dose treatment paradigm can produce effects that outlast acute drug exposure [8,9]. Ketamine has also been linked to striatal BDNF, ERK1/2, and mTOR signaling, altered dendritic spine morphology, changes in corticostriatal cross-frequency coupling, and suppression of LID-associated spectral activity [9,25,26]. Additional work suggests that ketamine’s anti-dyskinetic effects are pharmacologically complex and should not be attributed exclusively to NMDA receptor blockade or to a single basal ganglia pathway [10,29]. Lastly, others have done *in vivo* testing effects of serotonergic modulation in LID models using a dual pharmacological approach with quinpirole and SKF-81297 [30,31].

Together, these studies provide the rationale for testing whether ketamine reduces D2R-like or D1R-like receptor agonist-induced dyskinesia. The present study addressed this gap by testing whether co-administration of ketamine with quinpirole or SKF-81297 reduces LAO AIMs scores in L-DOPA-primed male Sprague Dawley rats. We hypothesized that ketamine will reduce quinpirole-induced but not SKF-81297-induced AIMs on the acute treatment day and that this reduction will persist. If supported, these findings would indicate that D2R-like receptor agonist-induced dyskinesia is sensitive to ketamine treatment and would support the possibility that indirect pathway-related mechanisms contribute to ketamine’s anti-dyskinetic effects.

## Material and Methods

**Table 1.**
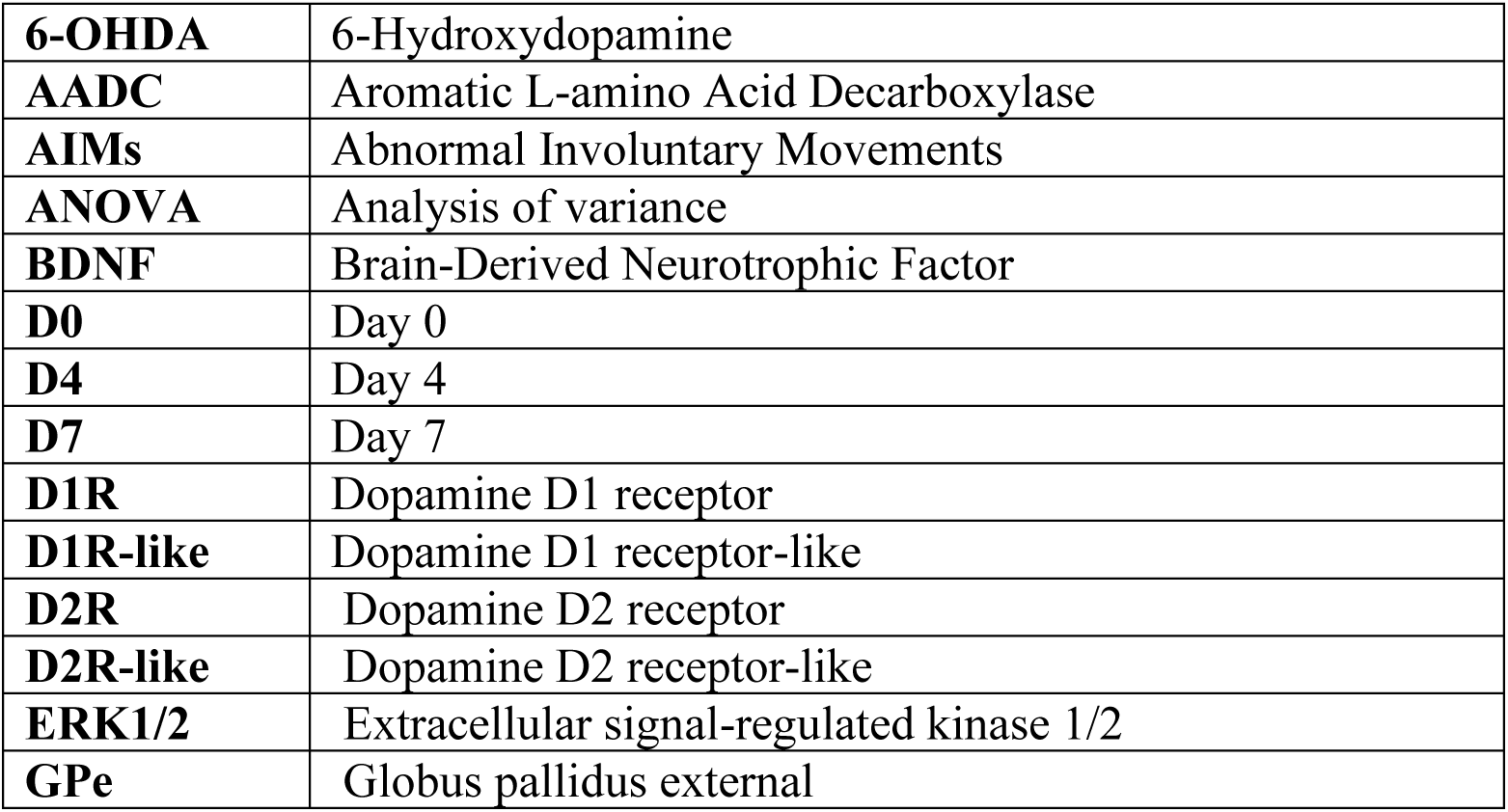
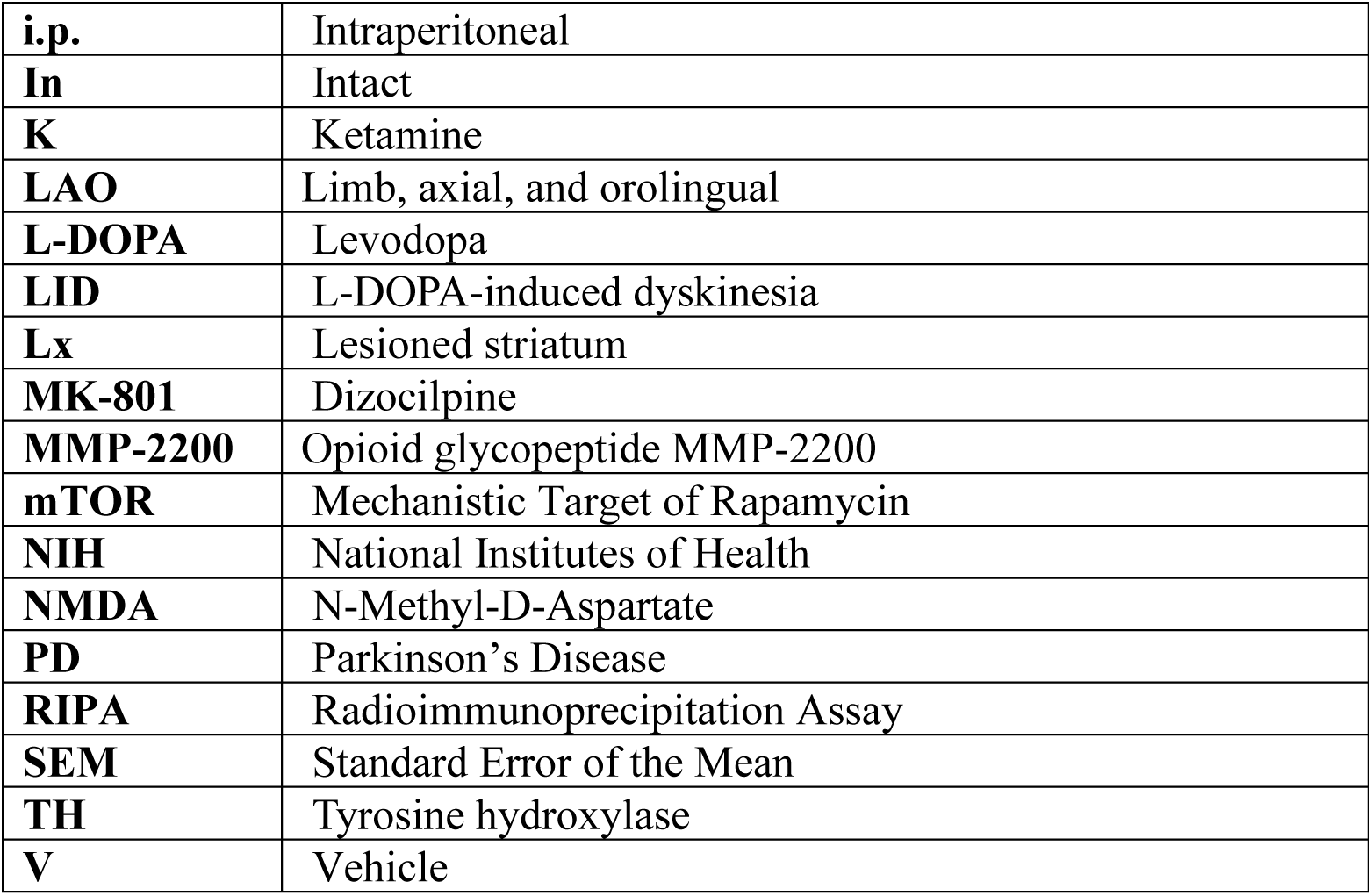
Abbreviations.

|  |  |
|---|---|
| **6-OHDA** | 6-Hydroxydopamine |
| **AADC** | Aromatic L-amino Acid Decarboxylase |
| **AIMs** | Abnormal Involuntary Movements |
| **ANOVA** | Analysis of variance |
| **BDNF** | Brain-Derived Neurotrophic Factor |
| **D0** | Day 0 |
| **D4** | Day 4 |
| **D7** | Day 7 |
| **D1R** | Dopamine D1 receptor |
| **D1R-like** | Dopamine D1 receptor-like |
| **D2R** | Dopamine D2 receptor |
| **D2R-like** | Dopamine D2 receptor-like |
| **ERK1/2** | Extracellular signal-regulated kinase 1/2 |
| **GPe** | Globus pallidus external |
| **i.p.** | Intraperitoneal |
| **In** | Intact |
| **K** | Ketamine |
| **LAO** | Limb, axial, and orolingual |
| **L-DOPA** | Levodopa |
| **LID** | L-DOPA-induced dyskinesia |
| **Lx** | Lesioned striatum |
| **MK-801** | Dizocilpine |
| **MMP-2200** | Opioid glycopeptide MMP-2200 |
| **mTOR** | Mechanistic Target of Rapamycin |
| **NIH** | National Institutes of Health |
| **NMDA** | N-Methyl-D-Aspartate |
| **PD** | Parkinson’s Disease |
| **RIPA** | Radioimmunoprecipitation Assay |
| **SEM** | Standard Error of the Mean |
| **TH** | Tyrosine hydroxylase |
| **V** | Vehicle |

### Animals

For Experiment 1 (D2R-like agonist study), a total of 18 adult male Sprague Dawley rats were included. Rats weighed 225 g at arrival and were obtained from Envigo RMC Inc. (Indianapolis, IN). Animals were housed two to three per cage throughout the study in a temperature- and humidity-controlled room on a 12-h reverse light/dark cycle. Food and water were provided *ad libitum*. Animals were assigned to one of two treatment groups: quinpirole + vehicle (n = 9) or quinpirole + ketamine (n = 9). For Experiment 2 (D1R-like agonist study), a separate cohort of 18 adult male Sprague Dawley rats was assigned to one of two treatment groups: SKF-81297 + vehicle (n = 9) or SKF-81297 + ketamine (n = 9). Animals were allowed to acclimate to the animal facility before the start of experimental procedures. Behavioral testing was performed during the animals’ active phase. All animal studies were approved by the Institutional Animal Care and Use Committee at The University of Arizona and were performed in accordance with the NIH Guidelines for the Care and Use of Laboratory Animals and the ARRIVE guidelines.

### Unilateral 6-hydroxydopamine (6-OHDA) lesion

Rats were pretreated 30 min prior to 6-OHDA administration with desipramine hydrochloride (12.5 mg/kg, i.p.) dissolved in sterile 0.9% normal saline containing 10% dimethyl sulfoxide (DMSO; Sigma-Aldrich, St. Louis, MO) to protect noradrenergic neurons. Animals were anesthetized with isoflurane (1.5–2.0%; VetOne, Boise, ID) delivered through a vaporizer (JD Medical, Phoenix, AZ) with 1.5 L oxygen/min and placed into a stereotaxic frame. A unilateral 6-OHDA lesion of the medial forebrain bundle was performed to induce nigrostriatal dopamine depletion.

6-OHDA hydrobromide (5.0 μg/μL in sterile 0.9% normal saline containing 0.02% ascorbic acid; Sigma-Aldrich) was administered at a rate of 0.5 μL/min using a microinjector (Stoelting Co., Wood Dale, IL) attached to a 26-gauge needle and 10 μL Hamilton syringe (Hamilton Co., Reno, NV). Two microliters of 6-OHDA were injected at each of two medial forebrain bundle coordinates, for a total of 10 μg 6-OHDA per coordinate. The stereotaxic coordinates were AP = −1.8 mm, ML = +2.0 mm, DV = −8.2 mm and AP = −2.8 mm, ML = +1.8 mm, DV = −8.2 mm, based on the Paxinos and Watson rat brain atlas. The needle was left in place for 5 min after each injection before removal to minimize backflow of the toxin. Given the established and standard use of this unilateral 6-OHDA lesion model in the field and in our laboratory, sham surgery groups were not included [7–15].

### Amphetamine-induced rotation test

Lesion severity was assessed using amphetamine-induced rotational behavior following recovery from unilateral 6-OHDA lesion surgery. This behavioral test was used to evaluate the degree of functional dopaminergic asymmetry produced by the lesion. In rats with unilateral nigrostriatal dopamine depletion, amphetamine induces dopamine release primarily from the intact hemisphere, producing rotational behavior biased toward the lesioned side. Therefore, increased net ipsilateral rotations provide an indirect behavioral measure of successful unilateral dopaminergic lesioning.

Rats received dextro-amphetamine (5 mg/kg, i.p.; Sigma-Aldrich, St. Louis, MO) prepared in sterile 0.9% saline and were placed individually into a clear Plexiglas cylinder measuring 38 cm in diameter × 38 cm in height. Rotational behavior was recorded for 1 minute every 5 minutes over a total testing period of 60 minutes. Both ipsilateral and contralateral rotations relative to the lesion side were recorded.

Net ipsilateral rotations were calculated by subtracting contralateral rotations from ipsilateral rotations and were expressed as rotations per minute. Rats were included in the study if they exhibited a net ipsilateral rotation score of ≥4 rotations/min, confirming sufficient unilateral dopaminergic lesion severity (>90%) for subsequent L-DOPA priming and dyskinesia assessment. All animals included in the final cohort met this inclusion criterion, resulting in a final sample size of n = 18 male Sprague Dawley rats.

### Drug and Vehicle Preparation

All systemic pharmacological agents were administered intraperitoneally at a standardized injection volume of 1 mL/kg body weight. L-DOPA (6 mg/kg, i.p.; Sigma-Aldrich) was dissolved in 0.9% USP-grade sterile saline and co-administered with benserazide hydrochloride (12 mg/kg, i.p.; Sigma-Aldrich), a peripherally acting aromatic L-amino acid decarboxylase inhibitor used to limit peripheral conversion of L-DOPA to dopamine and enhance central L-DOPA availability. Ketamine (20 mg/kg, i.p.; VetOne) was diluted in 0.9% USP-grade sterile saline. Quinpirole (0.2 mg/kg, i.p.), a D2R-like receptor agonist, and SKF-81297 (0.8 mg/kg, subcutaneous, s.c.), a D1R-like receptor agonist, was dissolved in 0.9% sterile saline and administered to induce D2R-like agonist-mediated AIMs, consistent with prior Falk laboratory studies using quinpirole-induced dyskinesia paradigms [7,15]. Vehicle-treated animals received 0.9% USP-grade sterile saline i.p. Drug solutions were prepared fresh on the day of experimentation and administered according to treatment group assignment.

### L-DOPA-induced dyskinesia (LID) rating scale

Dyskinesia severity was quantified by an investigator blinded to treatment group using the established AIM rating scale for the unilateral 6-OHDA rat model [11,13,14]. AIMs were scored as axial, limb, and orolingual movements. Axial AIMs included dystonic posturing of the neck and upper body; limb AIMs included abnormal movements of the forelimb contralateral to the lesion; and orolingual AIMs included vacuous jaw movements and tongue protrusions. Behavioral assessments were conducted for 1-minute observation periods every 20 minutes for 180 minutes following L-DOPA or quinpirole administration. Each AIM subtype was scored from 0 to 4, with 0 indicating no dyskinesia and 4 indicating continuous, non-interruptible dyskinetic movements. Limb, axial, and orolingual AIMs were summed to generate total LAO AIM scores, which were used as the primary measure of dyskinesia severity.

### Experimental design and treatment paradigm

This experiment (**Figure 1A**) was designed to determine whether ketamine reduces D2R-like receptor agonist-induced dyskinesia in male Sprague Dawley rats. Following unilateral 6-OHDA lesion and recovery, lesion severity was assessed using amphetamine-induced rotations. Animals were then primed with daily L-DOPA methyl ester (6 mg/kg, i.p.) and benserazide hydrochloride (12 mg/kg, i.p.) for 3 weeks to establish a dyskinetic state. After priming, L-DOPA-induced AIMs were maintained with twice-weekly L-DOPA administration, and AIM scores were reassessed every 4–7 days until a stable L-DOPA baseline was established.

**Figure 1.**
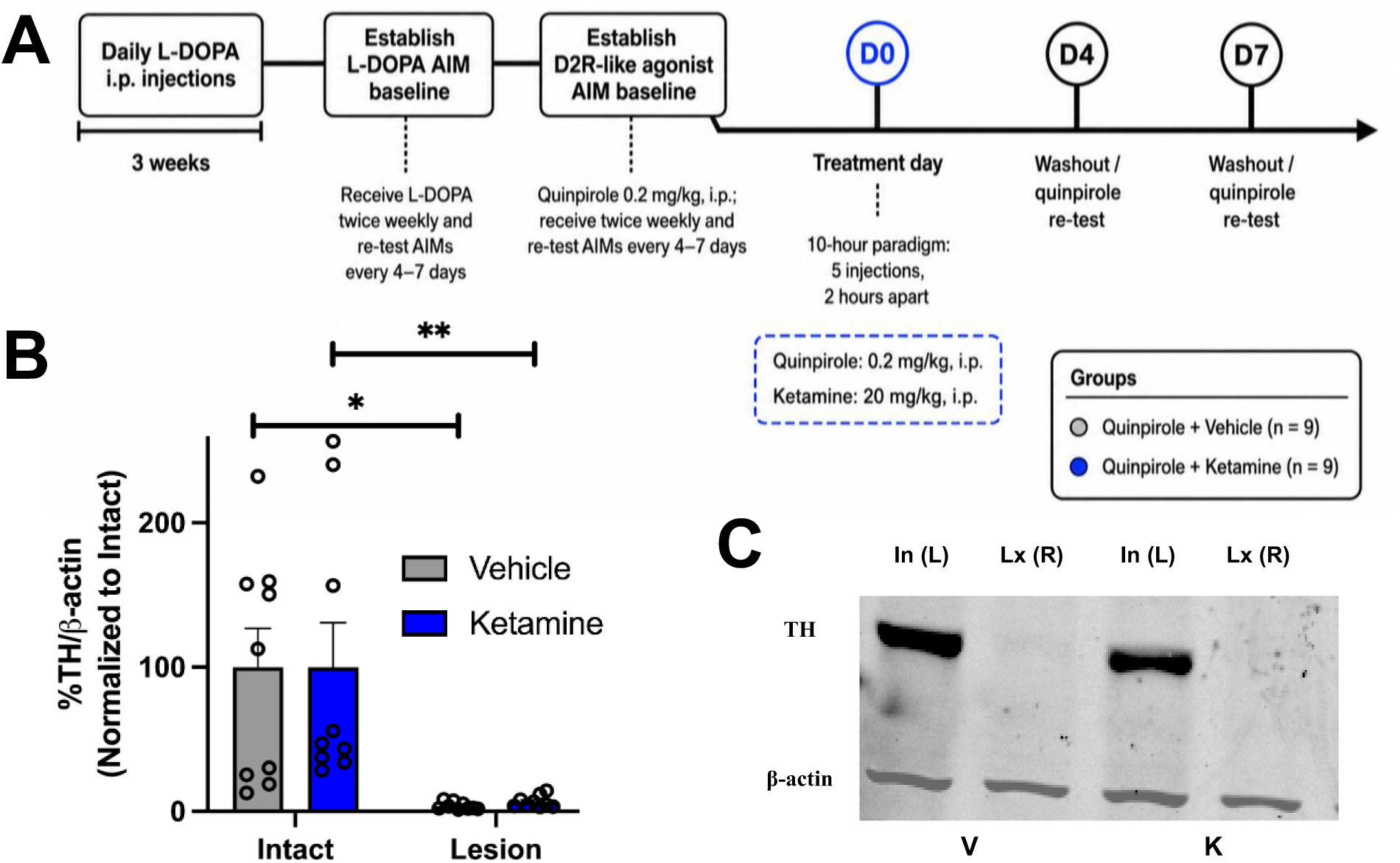
Study timeline and Verification of 6-ODHA lesion for D2 receptor-like study. **(A)** Male Sprague Dawley rats received a unilateral 6-OHDA lesion, recovered for 3 weeks, and then underwent amphetamine-induced rotations to assess lesion severity. Animals were then primed with daily L-DOPA for 3 weeks before L-DOPA AIM baselines and quinpirole-induced AIM baselines were established. Quinpirole (0.2 mg/kg, i.p.) was used to induce D2R-like agonist-induced AIMs (3 stable baselines were established). On D0, animals received quinpirole with either vehicle or ketamine (20 mg/kg, i.p.) using a 10-hour treatment paradigm consisting of 5 injections administered 2 hours apart. AIMs were re-tested after washout on D4 and D7 following quinpirole administration. Experimental groups included quinpirole + vehicle (n = 9) and quinpirole + ketamine (n = 9). **(B)** TH protein expression was measured in intact and lesioned striatal tissue and normalized to β-actin. TH expression was significantly reduced in the lesioned striatum compared with the intact striatum in both vehicle-treated animals, t(8) = 2.949, *p* = 0.0185, and ketamine-treated animals, t(8) = 3.554, *p* = 0.0075, as determined by two-tailed paired t-tests. Representative western blots show TH and β-actin expression in intact and lesioned samples from vehicle- and ketamine-treated animals. Data are presented as mean ± SEM. In, intact striatum; Lx, lesioned striatum; V, vehicle; K, ketamine. \**p* < 0.05, \*\**p* < 0.01. **(C)** Representative western blots showing TH and β-actin protein expression in intact and lesioned striatal samples from vehicle- and ketamine-treated animals. TH signal was reduced in lesioned samples compared with intact samples in both treatment groups, while β-actin served as the loading control. In, intact striatum; Lx, lesioned striatum; V, vehicle; K, ketamine.

Animals were then transitioned to a D2R-like receptor agonist-induced dyskinesia paradigm. Quinpirole (0.2 mg/kg, i.p.) was administered twice weekly, and AIM scores were reassessed every 4–7 days until a stable quinpirole-induced AIM baseline was established. A total of 18 animals were assigned to one of two treatment groups: quinpirole + vehicle (n = 9) or quinpirole + ketamine (n = 9). On D0, animals underwent a 10-hour treatment paradigm consisting of five treatment time points administered 2 hours apart. At each time point, animals received two intraperitoneal injections. The quinpirole + ketamine group received quinpirole (0.2 mg/kg, i.p.) and ketamine (20 mg/kg, i.p.), whereas the quinpirole + vehicle group received quinpirole (0.2 mg/kg, i.p.) and 0.9% sterile saline vehicle.

AIM scores were assessed on D0 to determine the acute effect of ketamine on quinpirole-induced dyskinesia. To determine whether this effect persisted beyond the acute treatment day, animals were re-challenged with quinpirole (0.2 mg/kg, i.p.) and retested after washout on D4 and D7. Limb, axial, and orolingual AIMs were summed to generate total LAO AIM scores, which were used as the primary behavioral measure of dyskinesia severity. Because this experiment did not directly manipulate indirect pathway neurons, reductions in quinpirole-induced AIMs were interpreted as evidence that D2R-like receptor-associated and/or indirect pathway-related mechanisms may contribute to ketamine’s anti-dyskinetic effects, rather than proof of a direct or exclusive indirect pathway mechanism.

### Western blots

Whole striatal tissue was collected from both hemispheres following euthanasia by CO₂ inhalation, with the right hemisphere designated as the lesioned side. Brains were rapidly removed, placed in a brain matrix, sectioned using razor blades, and striatal tissue was dissected on ice. Samples were flash-frozen and stored at −80°C until protein extraction. Tissue collection for Experiments 1 and 2 was performed 1 h following the final ketamine or vehicle injection. Striatal tissue was homogenized using a bead homogenizer in RIPA buffer supplemented with protease and phosphatase inhibitor mini tablets. Protein concentration was determined using the Pierce bicinchoninic acid (BCA) protein assay according to manufacturer instructions.

Protein samples were prepared at a concentration of 1 μg/μL, and 6.8 μg total protein was loaded per lane onto Any-kD Mini-PROTEAN TGX™ precast gels (Bio-Rad Laboratories, Hercules, CA). Proteins were transferred using a Trans-Blot Turbo Transfer System (Bio-Rad Laboratories). A Chameleon Duo prestained protein ladder (LI-COR, Lincoln, NE) was used for molecular weight determination. Membranes were blocked using LI-COR blocking buffer and incubated overnight at 4°C on a cold shaker with primary antibodies against tyrosine hydroxylase (TH; 1:2000; Millipore, Billerica, MA) and β-actin (1:2000; Sigma-Aldrich). Membranes were washed using TBST containing 0.1% Tween-20 and incubated for 1 h at room temperature with IRDye 800CW donkey anti-chicken secondary antibody (1:20,000; LI-COR) and IRDye 490CW donkey anti-goat secondary antibody (1:20,000; LI-COR). Blots were imaged using an Odyssey CLx imaging system (LI-COR) and analyzed using Empiria Studio Software (LI-COR). Protein expression was quantified using normalization to β-actin.

### Statistical Analyses

Statistical analyses were performed using GraphPad Prism version 10.6.1 software (GraphPad Software, La Jolla, CA). Data are presented as mean ± SEM, and statistical significance was set at p < 0.05. Total LAO AIM scores were used as the primary behavioral measure of dyskinesia severity and were calculated as the sum of limb, axial, and orolingual AIMs across the 180-min behavioral testing session. Following ketamine or vehicle treatment, total LAO AIM scores were compared between quinpirole + vehicle and quinpirole + ketamine groups at D0, D4, and D7 using two-tailed Mann–Whitney U tests. Because three planned day-specific comparisons were performed, p values were corrected across D0, D4, and D7 using the Holm–Bonferroni method.

Time-course LAO AIM scores on D0 and D7 were analyzed across the 180-min observation period using a mixed-effects model with restricted maximum likelihood estimation, with treatment group and time as factors. Šídák’s multiple comparisons post hoc tests were used to compare quinpirole + ketamine versus quinpirole + vehicle groups at individual time points following quinpirole administration. Statistical significance was indicated on graphs as follows: \**p* < 0.05, \*\**p* < 0.01, \*\*\**p* < 0.001, and \*\*\*\**p* < 0.0001.

Western blot quantification of TH protein expression was analyzed using two-tailed paired t tests. TH expression was normalized to β-actin, and normalized TH/β-actin values from the lesioned striatum were compared with matched intact striatal values within the same animals. Analyses were performed separately for quinpirole + vehicle and quinpirole + ketamine groups.

## Results

### Confirmation of 6-OHDA lesion severity for Experiment 1

Initial lesion severity was assessed using amphetamine-induced rotational behavior. Following dextro-amphetamine administration, animals exhibited an average of 7.70 ± 2.11 net ipsilateral rotations/min (n = 18), confirming successful unilateral lesion induction prior to experimental testing. Western blot analysis was used to verify dopaminergic lesion severity post-hoc by measuring TH protein expression in intact and lesioned striatal tissue. TH expression was normalized to β-actin and compared between paired intact and lesioned samples from the same animals. TH expression was significantly reduced in the lesioned striatum compared with the intact striatum in both treatment groups, confirming successful 6-OHDA lesioning. In vehicle-treated animals, TH expression was significantly reduced in the lesioned striatum compared with the intact striatum, t(8) = 2.949, *p* = 0.0185. In ketamine-treated animals, TH expression was also significantly reduced in the lesioned striatum compared with the intact striatum, t(8) = 3.554, *p* = 0.0075. Representative western blots further showed reduced TH signal in lesioned samples relative to intact samples, with β-actin used as the loading control (**Figure 1B–C**).

### Quinpirole induced stable D2R-like agonist-induced AIMs

The purpose of this experiment was to determine whether ketamine reduces dyskinesia induced by D2R-like receptor activation. After 6-OHDA lesioning, animals were primed with L-DOPA to establish a dyskinetic state and were then transitioned to a D2R-like receptor agonist-induced dyskinesia paradigm using quinpirole (**Figure 1A**). Quinpirole was administered at 0.2 mg/kg, i.p., and limb, axial, and orolingual AIMs were scored to quantify dyskinesia severity. Before ketamine or vehicle treatment, animals underwent repeated quinpirole baseline testing to confirm that quinpirole reliably induced AIMs. Across three baseline sessions, quinpirole produced consistent total LAO AIM scores, indicating that D2R-like agonist-induced AIMs were established before treatment assignment (**Figure 2A**).

**Figure 2.**
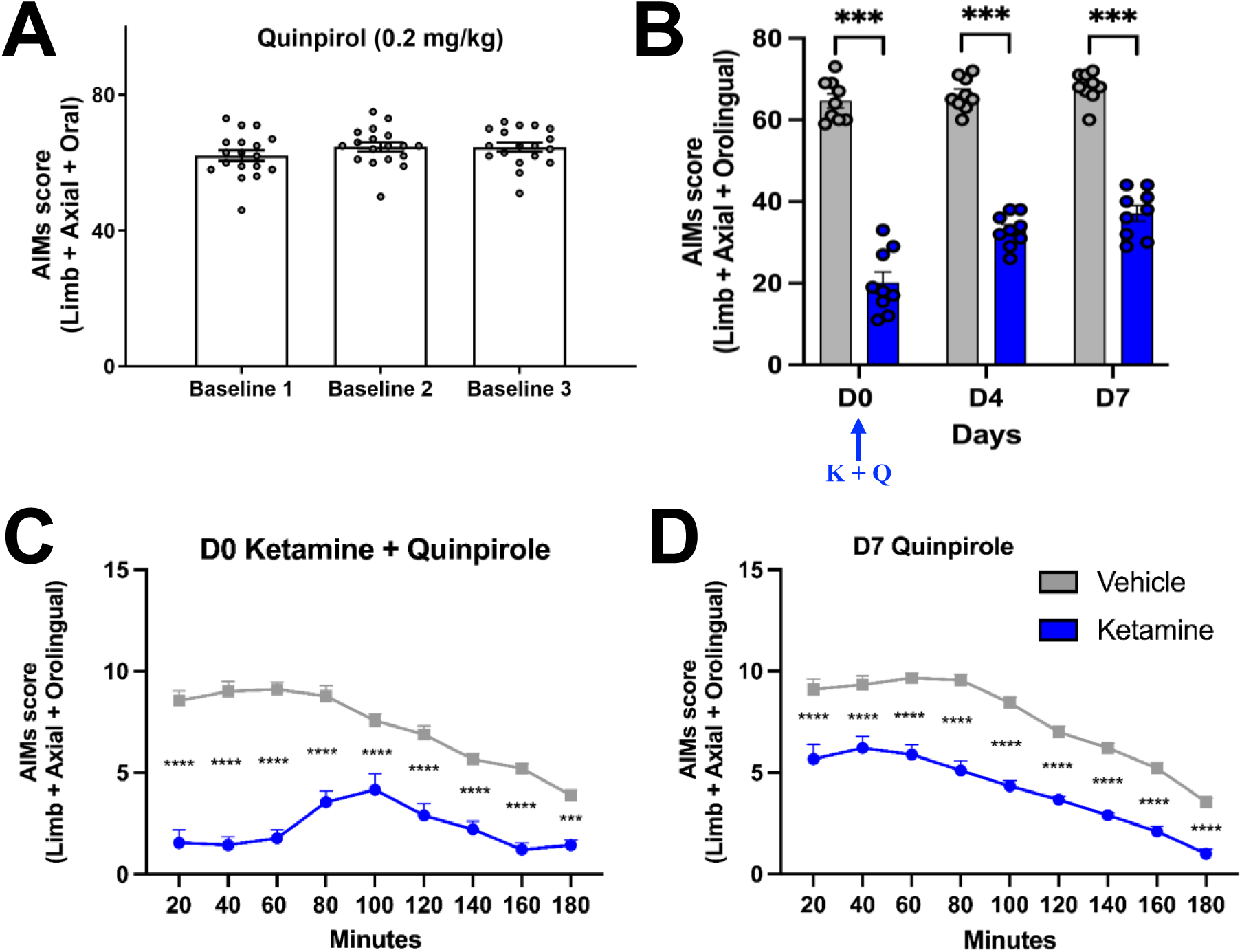
Ketamine reduced D2R-like agonist-induced AIMs in L-DOPA-primed male Sprague Dawley rats. **(A)** Quinpirole (0.2 mg/kg, i.p.) produced stable total LAO AIM scores across three baseline testing sessions, confirming establishment of D2R-like agonist-induced AIMs before treatment. **(B)** Ketamine significantly reduced quinpirole-induced total LAO AIM scores compared with vehicle on D0, D4, and D7. Day-specific comparisons were analyzed using two-tailed Mann–Whitney U tests with Holm–Bonferroni correction for multiple comparisons across the three testing days. Quinpirole + ketamine animals had significantly lower total LAO AIM scores than quinpirole + vehicle animals on D0, D4, and D7 (U = 0 for all comparisons; adjusted *p* = 0.00012 for all comparisons). LAO AIMs were calculated as the sum of limb, axial, and orolingual AIM scores; locomotor AIMs were not included. Data are presented as mean ± SEM. n = 9 animals/group. \*\*\**p* < 0.001. **(C)** On D0, ketamine significantly reduced quinpirole-induced AIM scores across the post-injection time course compared with vehicle. Mixed-effects analysis revealed significant effects of time [F(8,72) = 13.44, *p* < 0.0001], treatment [F(1,72) = 535.1, *p* < 0.0001], and time × treatment interaction [F(8,72) = 9.022, *p* < 0.0001]. Šídák’s multiple comparisons showed significant reductions at all time points from 20–180 min. **(D)** On D7, ketamine-treated animals continued to show reduced quinpirole-induced AIM scores across the post-quinpirole time course. Mixed-effects analysis revealed significant effects of time [F(8,72) = 47.93, p < 0.0001] and treatment [F(1,72) = 484.4, *p* < 0.0001], with no significant time × treatment interaction [F(8,72) = 1.446, p = 0.1924]. Šídák’s multiple comparisons showed significant reductions at all time points from 20–180 min. Data are presented as mean ± SEM. \*\**p* < 0.01, \*\*\**p* < 0.001, \*\*\*\**p* < 0.0001.

### Ketamine reduced total quinpirole-induced LAO AIM scores compared with vehicle

After stable quinpirole-induced AIMs were established, animals were assigned to quinpirole + vehicle or quinpirole + ketamine groups. On D0, animals underwent the 10-h treatment paradigm consisting of five treatment time points administered 2 h apart. Animals in the ketamine group received quinpirole (0.2 mg/kg, i.p.) and ketamine (20 mg/kg, i.p.), whereas animals in the vehicle group received quinpirole (0.2 mg/kg, i.p.) and 0.9% sterile saline. Ketamine significantly reduced quinpirole-induced total LAO AIM scores compared with vehicle across the post-treatment testing period (**Figure 2B**). Vehicle-treated animals maintained high AIM scores on D0, D4, and D7, whereas ketamine-treated animals showed markedly lower AIM scores at each testing day. Because AIM scores are ordinal behavioral ratings, day-specific comparisons were analyzed using nonparametric two-tailed Mann–Whitney U tests with Holm–Bonferroni correction across the three planned comparisons. Quinpirole + ketamine animals had significantly lower total LAO AIM scores than quinpirole + vehicle animals on D0 (U = 0, n = 9/group, adjusted *p* = 0.000123), D4 (U = 0, n = 9/group, adjusted p = 0.000123), and D7 (U = 0, n = 9/group, adjusted *p* = 0.000123). These findings demonstrate that ketamine reduced D2R-like receptor agonist-induced dyskinesia on the acute treatment day and that this reduction persisted after washout through D7.

### Ketamine acutely reduced quinpirole-induced AIM scores across the D0 time course

To further evaluate the acute effect of ketamine, AIM scores were analyzed across the post-quinpirole observation period on D0. Vehicle-treated animals showed robust quinpirole-induced AIMs across the time course, whereas ketamine-treated animals displayed lower AIM scores beginning at the earliest measured time point and continuing throughout the observation period (**Figure 2C**). Mixed-effects analysis revealed significant effects of time, F(8,72) = 13.44, *p* < 0.0001, treatment, F(1,72) = 535.1, *p* < 0.0001, and time × treatment interaction, F(8,72) = 9.022, *p* < 0.0001. Šídák’s multiple comparisons showed that ketamine significantly reduced quinpirole-induced AIM scores compared with vehicle at all measured time points from 20 to 180 minutes. These results indicate that ketamine did not reduce total AIM scores because of an isolated time point, but instead acutely suppressed quinpirole-induced dyskinesia across the post-injection time course.

### Ketamine-treated animals continued to show reduced quinpirole-induced AIMs on D7

To determine whether the effect of ketamine persisted beyond the acute treatment session, animals were re-challenged with quinpirole after washout. On D7, vehicle-treated animals continued to display robust quinpirole-induced AIMs across the post-quinpirole observation period (**Figure 2D**). In contrast, animals previously treated with quinpirole + ketamine showed lower AIM scores across the D7 time course. Mixed-effects analysis revealed significant effects of time, F(8,72) = 47.93, *p* < 0.0001, and treatment, F(1,72) = 484.4, p < 0.0001, with no significant time × treatment interaction, F(8,72) = 1.446, *p* = 0.1924. Šídák’s multiple comparisons showed that ketamine-treated animals had significantly lower AIM scores than vehicle-treated animals at all measured time points from 20 to 180 minutes. These findings indicate that ketamine-treated animals continued to show reduced quinpirole-induced dyskinesia after washout.

### Confirmation of 6-OHDA lesion severity for Experiment 2

The timeline for Experiment 2 is shown in **Figure 3A**. Initial lesion severity was assessed using amphetamine-induced rotational behavior. Following dextro-amphetamine administration, animals exhibited an average of 10.70 ± 0.46 net ipsilateral rotations/min (n = 18), confirming successful unilateral lesion induction prior to experimental testing. Western blot analysis was used to verify dopaminergic lesion severity post-hoc by measuring TH protein expression in intact and lesioned striatal tissue. TH expression was normalized to β-actin and compared between paired intact and lesioned samples from the same animals. TH expression was significantly reduced in the lesioned striatum compared with the intact striatum in both treatment groups, confirming successful 6-OHDA lesioning. In vehicle-treated animals, TH expression was significantly reduced in the lesioned striatum compared with the intact striatum, t(8) = 12.60, p < 0.0001. In ketamine-treated animals, TH expression was also significantly reduced in the lesioned striatum compared with the intact striatum, t(8) = 9.07, p < 0.0001. Representative western blots further showed reduced TH signal in lesioned samples relative to intact samples, with β-actin used as the loading control (**Figure 3B–C**).

**Figure 3.**
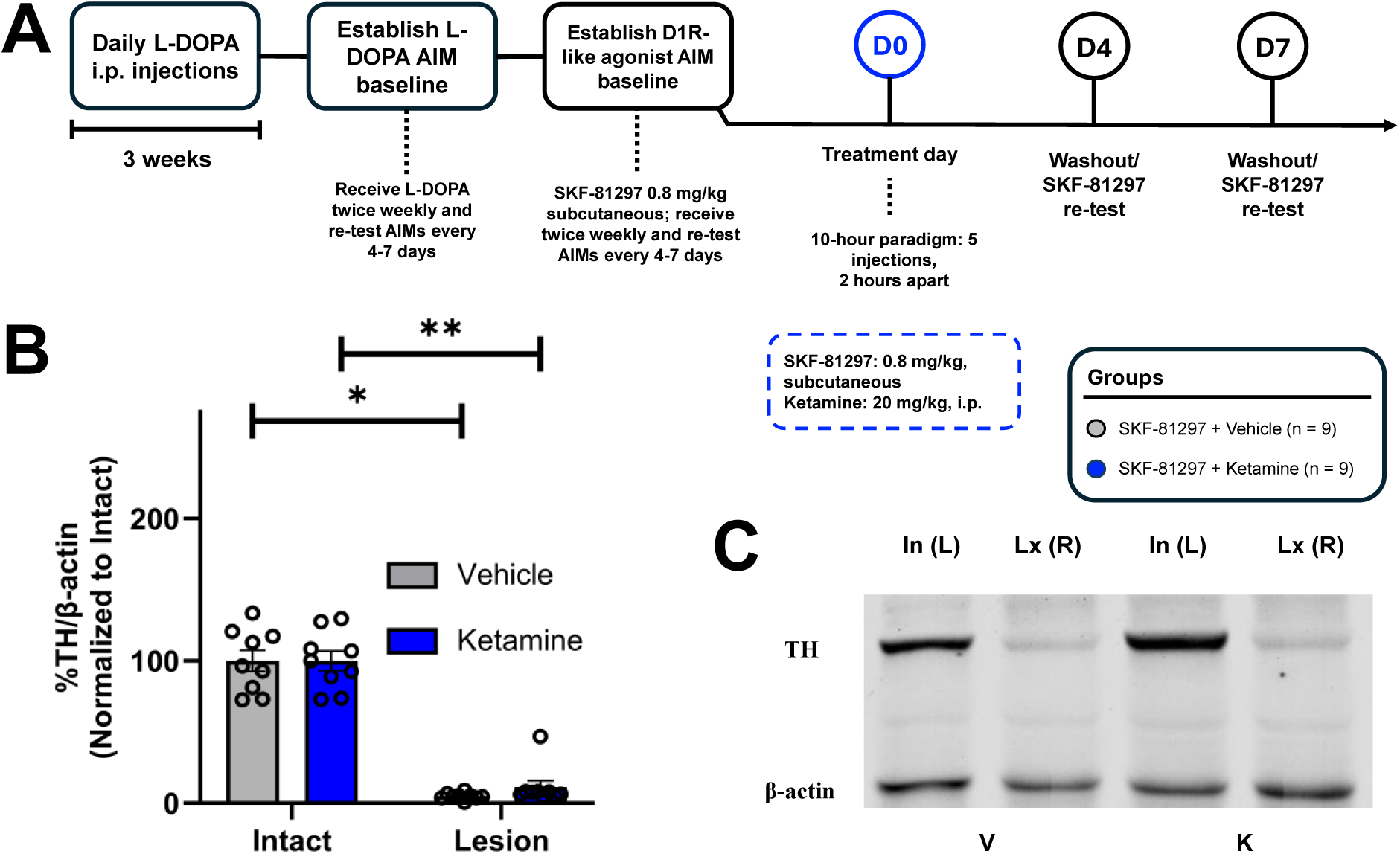
Study timeline and Verification of 6-OHDA lesion for D1 receptor-like study. **(A)** Male Sprague Dawley rats received a unilateral 6-OHDA lesion, recovered for 3 weeks, and then underwent amphetamine-induced rotations to assess lesion severity. Animals were then primed with daily L-DOPA for 3 weeks before L-DOPA AIM baselines and SKF-81297-induced AIM baselines were established. SKF-81297 (0.8 mg/kg; s.c.) was used to induce D1R-like agonist-induced AIMs (3 stable baselines were established). On D0, animals received SKF-81297 with either vehicle or ketamine (20 mg/kg, i.p.) using a 10-hour treatment paradigm consisting of 5 injections administered 2 hours apart. AIMs were re-tested after washout on D4 and D7 following SKF-81297 administration. Experimental groups included SKF-81297 + vehicle (n = 9) and SKF-81297 + ketamine (n = 9). **(B)** TH protein expression was measured in intact and lesioned striatal tissue and normalized to β-actin. TH expression was significantly reduced in the lesioned striatum compared with the intact striatum in both vehicle-treated animals, t(8) = 12.60, p < 0.0001, and ketamine-treated animals, t(8) = 9.07, p < 0.0001, as determined by two-tailed paired t-tests. Representative western blots show TH and β-actin expression in intact and lesioned samples from vehicle- and ketamine-treated animals. Data are presented as mean ± SEM. In, intact striatum; Lx, lesioned striatum; V, vehicle; K, ketamine. *p < 0.05, p < 0.01. **(C)** Representative western blots showing TH and β-actin protein expression in intact and lesioned striatal samples from vehicle- and ketamine-treated animals. TH signal was reduced in lesioned samples compared with intact samples in both treatment groups, while β-actin served as the loading control. In, intact striatum; Lx, lesioned striatum; V, vehicle; K, ketamine.

### SKF-81297 induced stable D1R-like agonist-induced AIMs

The purpose of this experiment was to determine whether ketamine reduces dyskinesia induced by D1R-like receptor activation. After 6-OHDA lesioning, animals were primed with L-DOPA to establish a dyskinetic state and were then transitioned to a D1R-like receptor agonist-induced dyskinesia paradigm using SKF-81297 (**Figure 3A**). SKF-81297 was administered at 0.8 mg/kg, s.c., and limb, axial, and orolingual AIMs were scored to quantify dyskinesia severity. Before ketamine or vehicle treatment, animals underwent repeated SKF-81297 baseline testing to confirm that SKF-81297 reliably induced AIMs. Across three baseline sessions, SKF-81297 produced consistent total LAO AIM scores, indicating that D1R-like agonist-induced AIMs were established before treatment assignment (**Figure 4A**).

**Figure 4.**
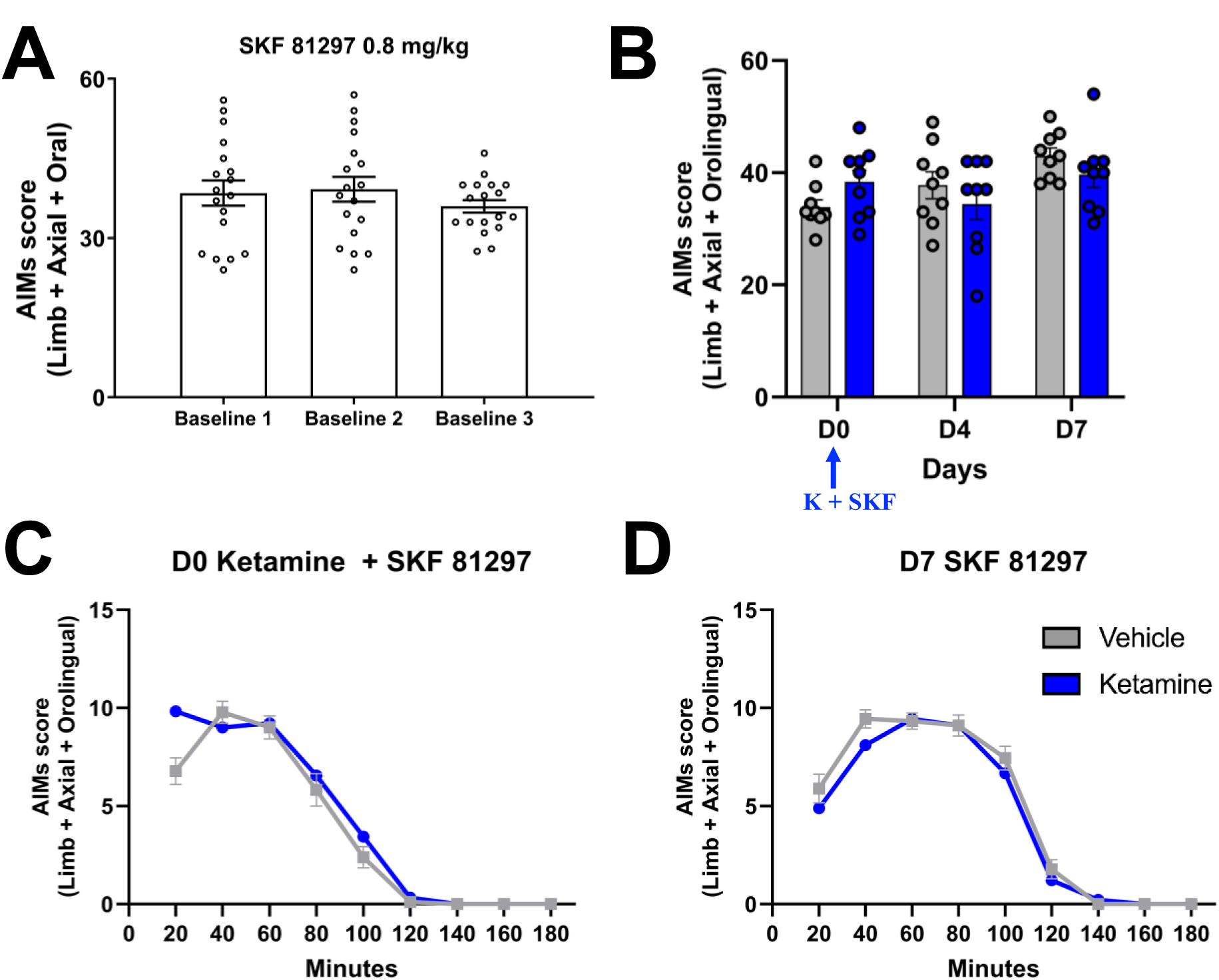
Ketamine did not alter D1R-like agonist-induced AIMs in L-DOPA-primed male Sprague Dawley rats. **(A)** SKF-81297 (0.8 mg/kg; s.c.) produced stable total LAO AIM scores across three baseline testing sessions, confirming establishment of D1R-like agonist-induced AIMs before treatment. **(B)** Ketamine did not significantly alter SKF-81297-induced total LAO AIM scores compared with vehicle on D0, D4, or D7. Day-specific comparisons were analyzed using two-tailed Mann–Whitney U tests with Holm–Bonferroni correction across the three testing days. SKF-81297 + ketamine animals showed no significant differences in total LAO AIM scores compared with SKF-81297 + vehicle animals on D0 (U = 23.50, n = 9/group, adjusted p = 0.4251), D4 (U = 34.00, n = 9/group, adjusted p = 0.6034), or D7 (U = 25.00, n = 9/group, adjusted p = 0.3628). LAO AIMs were calculated as the sum of limb, axial, and orolingual AIM scores; locomotor AIMs were not included. Data are presented as mean ± SEM. n = 9 animals/group. **(C)** On D0, ketamine did not significantly alter SKF-81297-induced AIM scores across the post-injection time course compared with vehicle. Mixed-effects analysis revealed a significant effect of time [F(8, 128) = 158.4, p < 0.0001], no significant main effect of treatment [F(1, 16) = 3.026, p = 0.1011], and a significant time × treatment interaction [F(8, 128) = 2.684, p = 0.0092]. Šídák’s multiple comparisons showed no significant differences at any time point from 20–180 min. **(D)** On D7, both vehicle- and ketamine-treated animals displayed comparable SKF-81297-induced AIM scores across the post-injection time course. Mixed-effects analysis revealed a significant effect of time [F(8, 128) = 172.5, p < 0.0001], with no significant main effect of treatment [F(1, 16) = 1.562, p = 0.2293] and no time × treatment interaction [F(8, 128) = 0.7786, p = 0.6222]. Šídák’s multiple comparisons showed no significant differences at any time point from 20–180 min. Data are presented as mean ± SEM.

### Ketamine did not significantly alter total SKF-81297-induced LAO AIM scores

After stable SKF-81297-induced AIMs were established, animals were assigned to SKF-81297 + vehicle or SKF-81297 + ketamine groups. On D0, animals underwent the 10-h treatment paradigm consisting of five treatment time points administered 2 h apart. Animals in the ketamine group received SKF-81297 (0.8 mg/kg, s.c.) and ketamine (20 mg/kg, i.p.), whereas animals in the vehicle group received SKF-81297 (0.8 mg/kg, s.c.) and 0.9% sterile saline. Ketamine did not significantly alter SKF-81297-induced total LAO AIM scores compared with vehicle across the post-treatment testing period (**Figure 4B**). Both vehicle- and ketamine-treated animals maintained high AIM scores on D0, D4, and D7. Because AIM scores are ordinal behavioral ratings, day-specific comparisons were analyzed using nonparametric two-tailed Mann–Whitney U tests with Holm–Bonferroni correction across the three planned comparisons. SKF-81297 + ketamine animals showed no significant differences in total LAO AIM scores compared with SKF-81297 + vehicle animals on D0 SKF-81297 + ketamine animals showed no significant differences in total LAO AIM scores compared with SKF-81297 + vehicle animals on D0 (U = 23.50, n = 9/group, adjusted p = 0.4251), D4 (U = 34.00, n = 9/group, adjusted p = 0.6034), or D7 (U = 25.00, n = 9/group, adjusted p = 0.3628). These findings demonstrate that ketamine did not reduce D1R-like receptor agonist-induced dyskinesia on the acute treatment day, nor after washout through D7. These findings demonstrate that ketamine did not reduce D1R-like receptor agonist-induced dyskinesia on the acute treatment day, nor after washout through D7.

### Ketamine did not alter SKF-81297-induced AIM scores across the D0 time course

To further evaluate the acute effect of ketamine, AIM scores were analyzed across the post-SKF-81297 observation period on D0. Vehicle-treated and ketamine-treated animals displayed comparable SKF-81297-induced AIMs across the time course (**Figure 4C**). Mixed-effects analysis revealed a significant effect of time (F(8, 128) = 158.4, p < 0.0001), no significant main effect of treatment (F(1, 16) = 3.026, p = 0.1011), and a significant time × treatment interaction (F(8, 128) = 2.684, p = 0.0092). Šídák’s multiple comparisons showed no significant differences in SKF-81297-induced AIM scores between ketamine- and vehicle-treated animals at any measured time point from 20 to 180 minutes. These results indicate that ketamine failed to acutely suppress D1R-like agonist-induced dyskinesia across the post-injection time course.

### Ketamine-treated animals had no significant reduction in SKF-81297-induced AIMs on D7

To determine whether any delayed effect of ketamine emerged after washout, animals were re-challenged with SKF-81297 on D7. On D7, both vehicle-treated and ketamine-treated animals continued to display robust SKF-81297-induced AIMs across the post-injection observation period (**Figure 4D**). Mixed-effects analysis revealed a significant effect of time (F(8, 128) = 172.5, p < 0.0001), with no significant main effect of treatment (F(1, 16) = 1.562, p = 0.2293), and no significant time × treatment interaction (F(8, 128) = 0.7786, p = 0.6222). Šídák’s multiple comparisons showed no significant differences between treatment groups at any measured time point from 20 to 180 minutes. These findings indicate that ketamine-treated animals showed no persistent reduction in D1R-like agonist-induced dyskinesia following washout.

## Discussion

The present study tested whether low-dose ketamine reduces dyskinesia induced by either D2R-like or D1R-like receptor activation in male Sprague Dawley rats previously rendered dyskinetic by L-DOPA priming. TH western blot verification confirmed dopaminergic denervation in both treatment groups (**Figure 1B–C**), and repeated quinpirole baseline testing showed stable D2R-like agonist-induced AIMs before ketamine or vehicle treatment (**Figure 2A**). The major finding was that ketamine significantly reduced quinpirole-induced LAO AIM scores compared with vehicle on D0, D4, and D7 (**Figure 2B**). Time-course analyses further showed that ketamine reduced AIM scores across the D0 observation period and that this reduction remained evident on D7 (**Figure 2C–D**). Together, these findings support the conclusion that ketamine can suppress dyskinesia induced by D2R-like receptor agonist challenge. Importantly, these data do not demonstrate that ketamine directly targets D2 receptors or acts exclusively through the indirect pathway. Rather, they suggest that D2R-like receptor-associated dyskinesia is sensitive to ketamine treatment. Crucially, when evaluating D1R-like receptor activation, ketamine failed to attenuate SKF-81297-induced LAO AIMs on the acute treatment day or following washout on D4 and D7 (**Figure 4**). This striking divergence demonstrates that ketamine does not globally dampen all dopamine receptor-driven dyskinesias, but instead exhibits clear selectivity for D2R-like, over D1R-like, agonist-induced motor abnormalities.

A critical first observation was that quinpirole produced stable AIM scores across baseline testing sessions before treatment assignment. This stability was important because D2R-like agonist-induced AIMs depend on prior dopaminergic priming history and are not necessarily equivalent to L-DOPA-induced or D1R agonist-induced AIMs. Prior studies have shown that the expression of AIMs after dopamine receptor agonist challenge is shaped by the type, dose, and duration of dopaminergic stimulation used during priming [32,33]. Drake and colleagues reported that quinpirole challenge produced different AIM profiles depending on prior dopaminergic stimulation in 6-OHDA-lesioned rats, supporting the idea that D2R-like agonist-induced behaviors reflect the response of an already sensitized basal ganglia circuit rather than acute receptor activation alone [33]. This is directly relevant to the present study because animals were first primed with L-DOPA before being transitioned to quinpirole baseline testing. Therefore, quinpirole-induced AIMs occurred within a dyskinetic striatal state established by prior L-DOPA exposure [32,33].

Historically, dyskinesia models have heavily emphasized D1R-expressing direct pathway hyperactivity as the primary driver of LID. However, early work established that L-DOPA-induced dyskinesia is accompanied by altered striatal gene expression affecting D2R-associated indirect pathway markers as well, a distinct D2-specific contributor to the dyskinetic state [11]. From a clinical perspective, dyskinesia is increasingly understood not merely as overactivity in a single pathway, but as a severe network imbalance between the direct (GO) and indirect (NOGO) striatal output pathways. While direct D1R stimulation has long been viewed as the predominant dyskinetogenic trigger, these findings reinforce that modulating D2R-like signaling or indirect pathway network dynamics remains a viable strategy to re-establish circuit equilibrium and attenuate dyskinesia expression.

The ability of quinpirole to induce AIMs after L-DOPA priming is also consistent with prior work using dopamine receptor agonists to probe receptor-linked contributions to dyskinesia. Delfino and colleagues showed that dopamine agonists with different receptor selectivity induce distinct dyskinetic profiles in parkinsonian rats, with D1/D5 stimulation producing stronger dyskinesia and D2-family stimulation producing milder but measurable dyskinetic behavior under primed conditions [16]. Monville and colleagues further supported the use of selective dopamine receptor agonists and antagonists to examine receptor subtype contributions in the 6-OHDA model [34]. Together, these studies support the use of quinpirole-induced AIMs as a behavioral readout of D2R-like receptor-associated dyskinesia, while also reinforcing that systemic agonist challenge does not isolate one neuronal pathway with complete specificity [16,30,31].

The present findings extend our prior work showing that D2R-like agonist-induced dyskinesia is pharmacologically modifiable. Flores et al. reported that the opioid glycopeptide MMP-2200 reduced quinpirole-induced LAO AIMs by approximately 40% and quinpirole-induced locomotor AIMs by approximately 80% [7]. That study is especially relevant because it used quinpirole at 0.2 mg/kg, i.p., the same dose used in the present study, and demonstrated that quinpirole-induced dyskinesia can be reduced by a non-dopaminergic pharmacological intervention [7]. Flores et al. systematically evaluated both D1R- and D2R-driven dyskinesia paradigms, showing that MMP-2200 reduced quinpirole-induced LAO AIMs as well as D1R-mediated behaviors, demonstrating that both pathways contribute meaningfully to LID expression. The present results build upon this concept by demonstrating that while both receptor families drive dyskinetic output in primed animals, NMDA receptor-targeted intervention with sub-anesthetic ketamine displays clear pathway-specific selectivity for D2R-like over D1R-like activation. The present results are consistent with this prior work by showing that ketamine also suppresses quinpirole-induced AIMs. However, ketamine and MMP-2200 should not be interpreted as acting through the same mechanism, yet there could be overlap. MMP-2200 is a mixed mu/delta opioid glycopeptide agonist, and ketamine is a multifunctional compound best known as a non-competitive NMDA receptor antagonist, but is also a known PAM for opioid receptors [28], and has additional effects on glutamatergic, neuromodulatory, and plasticity-linked signaling systems [7,9,15]. Yet our prior work had shown that the acute anti-dyskinetic activity of ketamine cannot be blocked by the pan-opioid receptor antagonist naloxone [10]. That work rather pointed to the opioid activity of ketamine being involved in the acute anti-parkinsonian activity of ketamine, presumably mainly via the delta opioid activity.

The present results also relate to prior work with MK-801. Flores et al. showed that MK-801 reduced D2R-like agonist-induced LAO AIMs after quinpirole challenge but also induced ipsiversive rotations, suggesting a pro-parkinsonian behavioral effect in that model [15]. Paquette and colleagues similarly reported that MK-801 suppressed L-DOPA-induced AIMs only at doses that worsened parkinsonism [23]. These findings indicate that reducing AIM scores alone is not sufficient to establish therapeutic potential, particularly for strong noncompetitive NMDA receptor antagonists [23]. The present findings are important in this context because ketamine reduced quinpirole-induced AIMs and has previously shown sustained anti-dyskinetic and acute anti-parkinsonian effects in preclinical LID models [8,9]. However, additional motor assessments would be needed to determine whether ketamine reduces quinpirole-induced AIMs without impairing general motor function during D2R-like agonist challenge.

One of the most important aspects of this study was the persistence of ketamine’s effect for one week until the end of the experiment, tested on two days, D4 and D7. If ketamine reduced AIMs only through acute sedation, transient motor suppression, or short-lived interference with quinpirole responsiveness, the reduction would be expected to disappear after the D0 treatment period. Instead, ketamine-treated animals continued to show lower quinpirole-induced AIM scores after washout. This pattern is consistent with earlier studies by our group showing that sub-anesthetic ketamine produces long-lasting reductions in LID in 6-OHDA-lesioned rats [7,9,27,29]. Bartlett et al. first reported that a 10-hour sub-anesthetic ketamine paradigm reduced L-DOPA-induced AIMs beyond the acute exposure period, and later work showed that ketamine attenuates LID development and engages plasticity-related signaling pathways [8,9].

The sustained reduction in quinpirole-induced AIMs may reflect ketamine-induced changes in dyskinesia-related plasticity, although the present study did not directly test the molecular or circuit mechanisms responsible. LID has been linked to maladaptive striatal plasticity, altered corticostriatal synaptic function, and abnormal intracellular signaling [20,21]. Cenci and Konradi described LID as a disorder involving maladaptive striatal plasticity, while Picconi and colleagues reported loss of bidirectional corticostriatal synaptic plasticity in dyskinetic animals [21,22]. These concepts are relevant because ketamine’s effects on D4 and D7 suggest a behavioral change that outlasts acute drug exposure and may involve neural processes that maintain dyskinesia expression [8,9, 27,29].

Mechanistic work from our group further supports the possibility that ketamine modifies dyskinesia-related plasticity [36]. Bartlett et al. reported that sub-anesthetic ketamine attenuated LID development and implicated striatal BDNF release followed by ERK1/2 and mTOR signaling in its long-term anti-dyskinetic effects [9]. That study also linked ketamine treatment to changes in striatal dendritic spine morphology [9] and pointing to modulation of TrkB signaling [29]. Electrophysiological studies showed that low-dose ketamine altered corticostriatal cross-frequency coupling and suppressed LID-associated spectral activity, suggesting that ketamine can modify dyskinesia-related network activity [25,26,35].

The current findings should also be considered in the context of emerging evidence that D2R-expressing striatal neurons and indirect pathway-related circuits contribute to dyskinesia. Although D1 receptor-expressing direct pathway neurons have often been emphasized in LID, newer studies indicate that D2R-expressing neurons and downstream pallidal circuits also influence dyskinetic behavior [17,18]. Liu et al. reported that abnormal activity of D2R-expressing striatal neurons and downstream GPe neurons contributes to dyskinetic symptoms, while Sáez et al. showed that D2-type dopamine receptors and the striatopallidal pathway modulate LID [17,18]. Studies using more direct pathway manipulation have also shown that indirect pathway activity can influence dyskinesia expression [19,20]. Castela and colleagues reported that selective activation of the striatal indirect pathway suppressed LID, and Florio and colleagues showed that D2R signaling in striatal spiny neurons modulates dyskinesia severity [19,20]. These studies support the relevance of examining D2R-like agonist-induced AIMs, but they also highlight the limits of the present pharmacological approach.

Because quinpirole was administered systemically, the behavioral effects observed here likely reflect network-level responses rather than isolated activation of one cell type or pathway. Quinpirole activates D2R-like receptors, including D2 and D3 receptors, and does not selectively manipulate only indirect pathway spiny projection neurons. Therefore, the current results should be interpreted as evidence that ketamine reduces dyskinesia induced by D2R-like receptor activation, not as proof that ketamine directly acts on D2 receptors or exclusively corrects indirect pathway dysfunction. This distinction is important because ketamine has multiple pharmacological actions, and the present experiment did not include pathway-selective manipulations, receptor-specific blockade, electrophysiology, or molecular measures. Furthermore, these findings suggest that for NMDA receptor-targeted and ketamine-based therapeutic strategies, D2R-like or indirect pathway-associated circuitry represents the most promising pathway in terms of pharmacological responsiveness. Rather than blunting direct pathway activity, which could risk suppressing necessary pro-motor D1R signaling, sub-anesthetic ketamine preferentially modulates the hyperactive or maladaptive signaling associated with D2R-like receptor activation. This selective engagement provides a therapeutic window to suppress dyskinesia driven by D2R-like stimulation while sparing D1R-mediated motor behaviors. An important next step will be determining how ketamine’s anti-dyskinetic effects relate to direct and indirect pathway-associated dyskinesia mechanisms. Dopamine receptor agonist-induced AIM paradigms provide a useful framework for addressing this question because D1R- and D2R-family receptor activation engage distinct, though partially overlapping, circuit-level processes. The present study focused specifically on D2R-like receptor-induced dyskinesia; future work systematically comparing ketamine effects across D1R- and D2R-driven paradigms may help clarify whether ketamine preferentially influences particular dopamine receptor-associated circuitry or instead acts through convergent downstream mechanisms shared across dyskinetic states.

One limitation of this work is that the present experiment was conducted only in male Sprague Dawley rats. Therefore, future studies are needed to determine whether ketamine similarly reduces D2R-like agonist-induced dyskinesia in female animals. In addition, because ketamine has multiple pharmacological actions and this study did not directly manipulate indirect or direct pathway neurons, these results do not establish an exclusive indirect pathway mechanism. In addition, we could not provide structural validation of direct vs. indirect pathway involvement in our studies, and test if ketamine reduces excess mushroom spines specifically in identified D2R MSNs, since a Golgi stain would not allow for co-stain to distinguish D1R vs D2R-positive neurons. However, by systematically evaluating both D2R-like (quinpirole) and D1R-like (SKF-81297) agonist challenges, our findings demonstrate that sub-anesthetic ketamine selectively suppresses D2R-like, but not D1R-like, dopamine receptor-induced dyskinesia, supporting a preferentially D2R-associated or indirect pathway-related mechanism. Our studies were conducted in animals that were first primed with L-DOPA to mimic the clinical situation better, but future work could also utilize protocols to instead prime rat PD models with either D1R- or D2R-agonists [37,38].

In conclusion, sub-anesthetic ketamine selectively reduced D2R-like agonist-induced dyskinesia without altering D1R-like agonist-induced dyskinesia in L-DOPA-primed rats. Ketamine significantly decreased quinpirole-induced LAO AIMs on the acute treatment day, an effect that persisted through one-week post-treatment, whereas SKF-81297-induced AIMs remained entirely unaffected. These findings extend prior evidence that ketamine suppresses LID [8,9] by demonstrating efficacy in a D2R-like agonist-induced but not a D1R-like agonist AIM paradigm. This clear selectivity indicates that indirect (NOGO) pathway-associated processes, rather than direct (GO) pathway-driven mechanisms, contribute to ketamine’s anti-dyskinetic effects. This is clinically relevant because many currently used dopamine agonists primarily target D2-like receptors, whereas D1R-like agonists have been limited by adverse neuropsychiatric effects. Together, this work further supports the potential repurposing of sub-anesthetic ketamine as an adjunctive therapeutic strategy for individuals with LID.

## CRediT authorship contribution statement

**Raveena Parmar**: Formal analysis, Investigation, Methodology, Validation, Writing – original draft. **Jasmine R. Meredith:** Formal analysis, Investigation, Methodology, Validation, Writing – original draft. **Mitchell J. Bartlett**: Formal analysis, Investigation, Methodology, Supervision, Validation. **Emma G.E. Smith**: Investigation, Methodology. **Jeanelle M. Castro**: Investigation, Methodology. **Scott J. Sherman**: Writing – review & editing. **Gaurav Chattree**: Writing – review & editing. **Kathy Steece-Collier**: Writing – review & editing. **Torsten Falk**: Conceptualization, Funding acquisition, Resources, Supervision, Writing – review & editing.

## Acknowledgements

Research support: National Institute of Health NINDS [R01 NS122805, TF], Training grant funding NIH T35 HL007479 [MJB], NIH NINDS Diversity Supplement R01 NS122805-03S1 [RP], and the Mary Mack Estate Gift (SJS, TF).

